# An ultra-high-throughput method for measuring biomolecular activities

**DOI:** 10.1101/2022.03.09.483646

**Authors:** Boqiang Tu, Vikram Sundar, Kevin M. Esvelt

## Abstract

Large datasets of biomolecular activities are crucial for protein engineering, yet their scarcity due to limited experimental throughput hampers progress. We introduce Direct High-throughput Activity Recording and Measurement Assay (DHARMA), an innovative method enabling ultra-high-throughput measurement of biomolecular activities. DHARMA employs molecular recording techniques to link activity directly to editing rates of DNA segments contiguous with the coding sequence of biomolecule of interest. Leveraging a Bayesian inference-based denoising model, we mapped the fitness landscape of TEV protease across 160,000 variants. Using these datasets, we benchmarked popular protein models and showed the impact of data size on model performance. We also developed circuit self-optimization strategies and demonstrated DHARMA’s capability to measure a wide range of biomolecular activities. DHARMA represents a leap forward, offering the machine learning community unparalleled datasets for accurate protein fitness prediction and enhancing our understanding of sequence-to-function relationships.

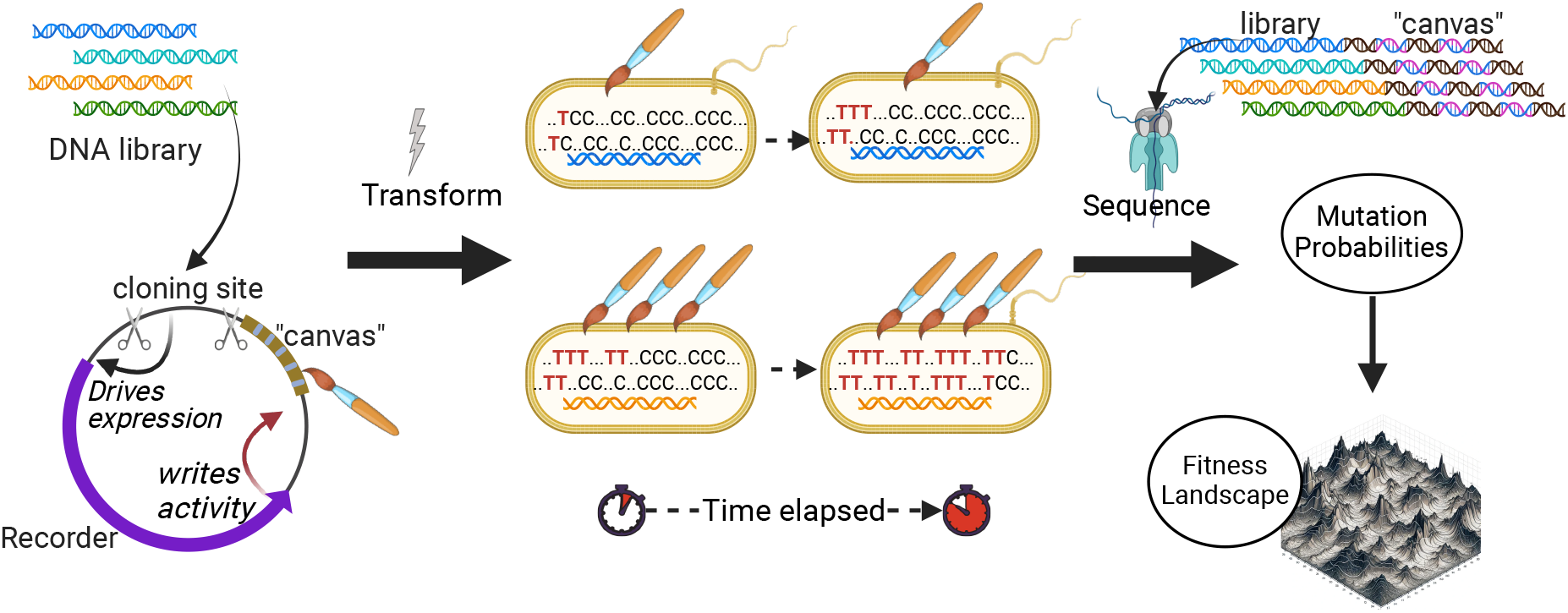

## Main

The field of protein engineering has experienced significant advancements over the past decades. From improving catalytic efficiency to modifying protein-protein interactions, powerful techniques based on directed evolution and rational design have fundamentally changed the landscape of protein engineering. While our evolving understanding of protein structures can enable effective targeted strategies for designing proteins with novel or improved properties, predicting protein fitness directly from amino acid sequences still remains a significant challenge. Unlike structure-focused protein machine learning based on sequence data, fitness prediction often requires functional data that are currently generated through laborious experimental techniques based on low-throughput biochemical and biophysical methods, typically *in vitro* enzyme activity and protein binding assays. These methods are unsuitable for generating large-scale datasets for training protein fitness machine learning (ML) models.

While some high-throughput methods exist for inferring fitness of protein variants, these methods, namely phage display, yeast display, and fluorescence-activated cell sorting (FACS), are often enrichment-based [1, 2, 3] and can introduce considerable noise into the data. Recently, a microfluidic-based high-throughput system for measuring enzyme activities have been developed [4], yet its throughput remains limited by the feature size of the chip, as each protein variant must be placed separately.

To effectively train protein fitness prediction models, a substantial amount of data is likely required, yet generating such large datasets using current methodologies is fraught with difficulties. These limitations highlight the urgent need for a novel approach that can provide high throughput, high accuracy, and broad accessibility, which could empower a community-based effort to produce the extensive datasets vital for advancing protein engineering research.

Here we present Direct High-throughput Activity Recording and Measurement Assay (DHARMA), a ultra-high-throughput method for measuring the activity of each member of a genetically encoded library over a wide dynamic range using only Nanopore sequencing. Unlike previous molecular recording methods that record external stimuli at the population level [5], DHARMA can measure the biomolecular activity of a given variant in each individual cell within a mixed population. Our implementation utilizes a base editor as a reporter protein to record the activity of a functional sequence onto a DNA “canvas” contiguous to each library member. The activity information recorded in the canvas and the identity of the corresponding variant are then simultaneously retrieved via nanopore sequencing. Each identified variant is then assigned a string of C to T mutations accumulated in the canvas. We tested the feasibility of the system using a set of well characterized promoters, performed optimized assays on large libraries of T7 RNA polymerase and TEV protease variants, and applied FLIGHTED, a custom machine learning method to remove experimental noise from the high-throughput data for subsequent predictive modeling.

## Results

### Quantitative measurement of biological activities using molecular recording

We began by generating plasmid constructs consisting of a base editor expressed under the control of a constitutive promoter, a canvas with multiple target sites for activity recording, and a GFP or luxAB expressed under the control of the same promoter to serve as a signal reference for relative activity quantification (Figure 1A). In the presence of sgRNA, such constructs simultaneously measure the promoter strength using both sequencing and either fluorescence or luminescence intensity. To challenge the system with a wide range of activity levels, we cloned a set of previously characterized insulated promoters and showed that Nanopore sequencing alone is sufficient for activity quantification and is well-calibrated (Figure 1B). Intriguingly, DHARMA offers a greater dynamic range than is achievable with fluorescence or luminescence due its temporal flexibility: sequencing early in the experiment can better distinguish between high-activity constructs when they have generated the optimal number of mutations, while sequencing later in the experiment provides low-activity variants with enough time to accumulate the same number (Figure 1C).

**Figure 1:**
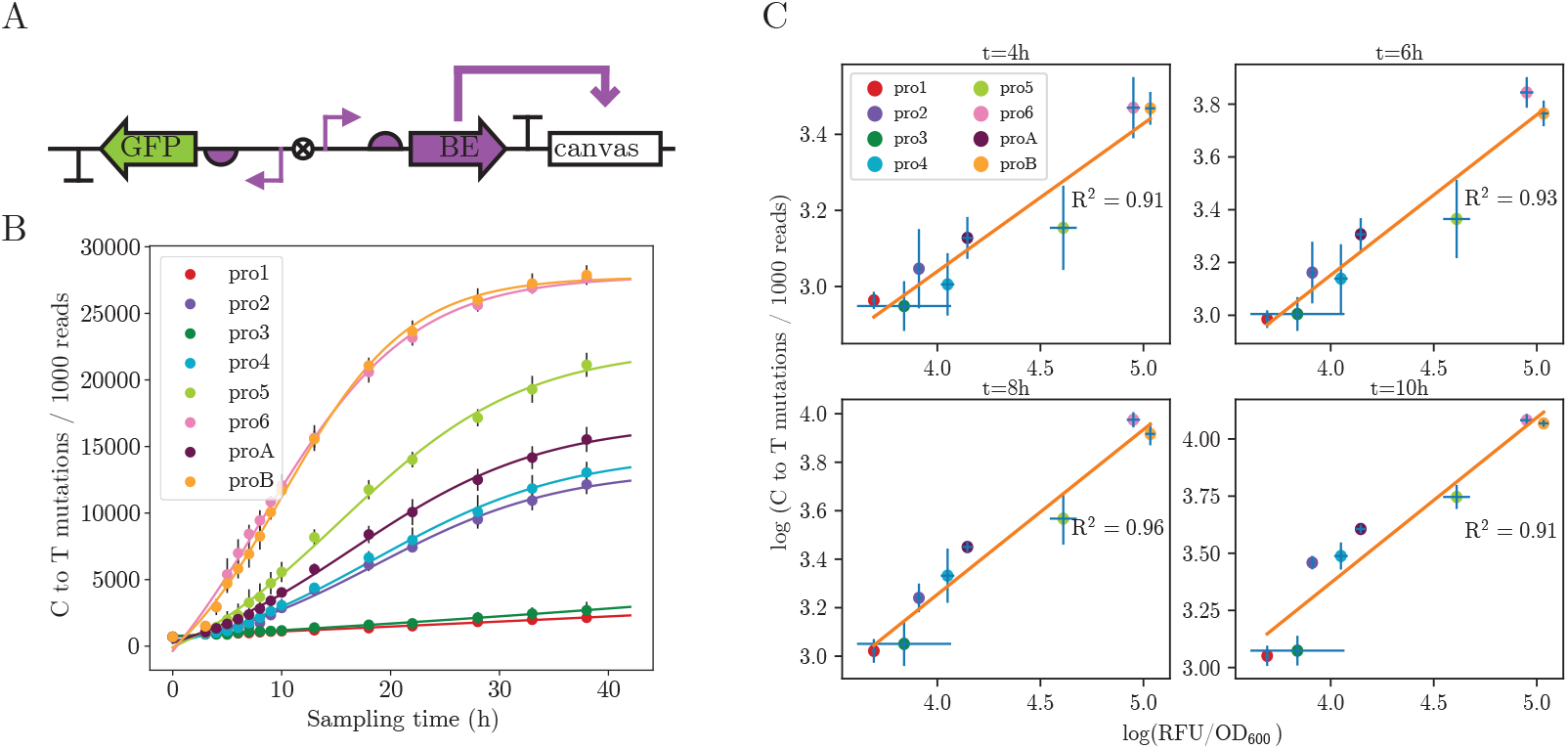
Molecular recording enables accurate activity measurement over a wide dynamic range. A. Diagram of promoter library constructs for validating molecular recording-based activity assay. Promoters colored in purple are the promoters of interest B. C-to-T mutation counts in the canvas region of each promoter construct fitted to sigmoidal functions. C. Correlation between the number of C-to-T mutations in the canvas region and the fluorescence intensity of GFP at different time points.

While coupling the activity of interest to the expression level of base editor in the context of promoter libraries yielded results that allow for direct and accurate quantification of biological activities, DHARMA assays targeting protein libraries require additional circuit optimization to achieve satisfactory dynamic range and signal-to-noise ratio as the circuits become more complex. To this end, we developed systematic circuit optimization strategies (Figure 2A) that leverage the ease of DHARMA library generation and the high-throughput nature of sequencing-based assays to essentially optimize DHARMA with DHARMA. After the initial library design that links the protein fitness or activity of interest to the expression of a mutagenic protein (e.g. the transcription, translation or post-translational modification of a base editor), we use the Golden Gate method to combinatorially assemble a library of plasmid constructs with varying regulatory components and coding sequences (including wild-type and truncated versions), which likely contain candidates that could potentially capture the full range of activities of the protein library. As shown in the example (Figure 2B), thousands of circuit designs can be assembled and tested in a single experiment. We then directly use DHARMA to characterize the performance of each individual member of the construct library by transforming the library into cells later grown in a continuous culture and sampled periodically for sequencing, under conditions identical to those used for the final library. Using long read sequencing, we retrieve both the identities of the relevant components and the corresponding activity at a given sampling time. This allows us to assess the dynamic range of each construct design spanning all timepoints in high throughput. The sequencing data is then fed into an alignment and demultiplexing pipeline to simultaneously identify mutations in the coding sequencing and their corresponding activities. The fitness data are then used to estimate the dynamic range and noise level of each construct design. If the top candidates in the construct library do not yield a dynamic range that allow us to accurately measure individual activities with less than 100 raw reads, we can use the data to inform the next round of construct library design. This design-build-test cycle can be iterated until the desired dynamic range and signal-to-noise ratio are achieved. Using this optimization strategy, we successfully designed and optimized biological circuits to measure a wide range of activities, including those of T7 RNA polymerase and TEV protease.

**Figure 2:**
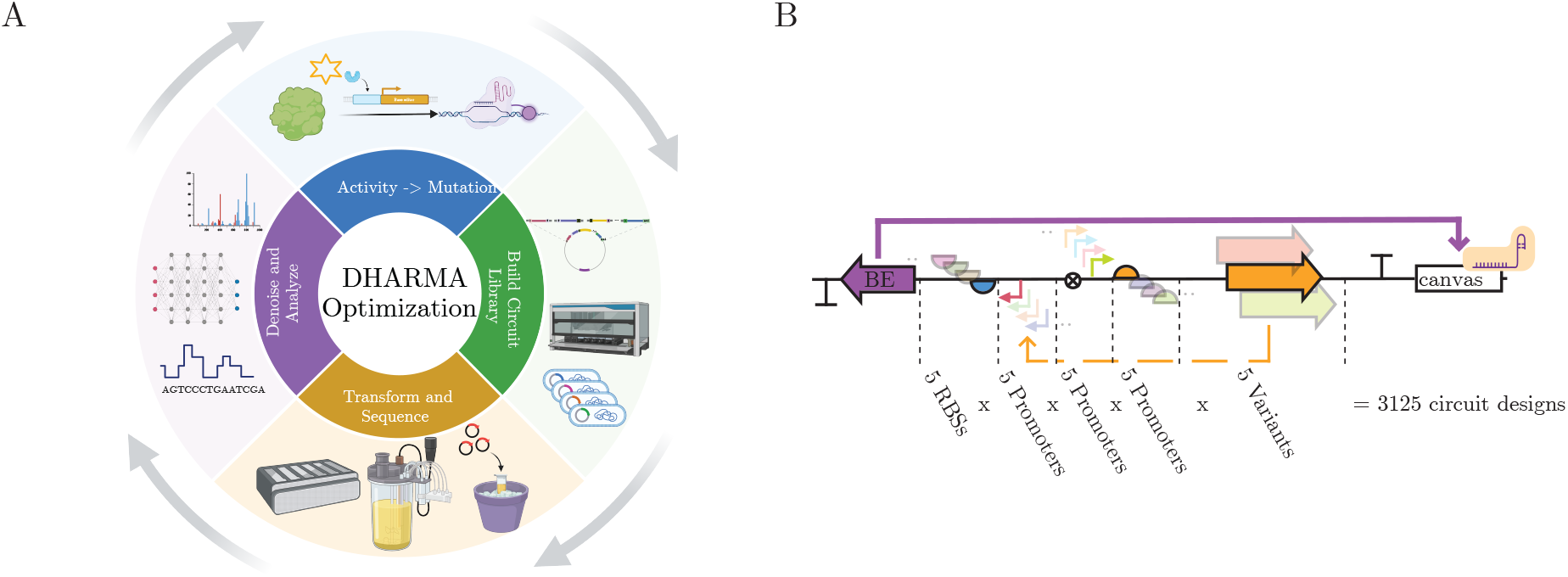
Optimization strategies for DHARMA circuits. A. General optimization workflow for DHARMA circuits. B. Example of a combinatorial library of DHARMA optimization constructs. Each unique combination is associated with a canvas region where corresponding activity is recorded.

### High-throughput activity quantification of T7 RNA polymerase variants with error correction

Next, we tested the feasibility of measuring the activity of each member of a small library by performing site-saturation mutagenesis on three residues of the T7 RNA polymerase specificity loop and constructing a circuit in which T7 RNAP variants differentially drive the expression of a base editor under the control of a T3 promoter (Figure 3A). After transforming cells with constructed library of T7 RNAP and subjected them to nanopore sequencing, we generated a set of raw reads each of which contain both variant identity and corresponding canvas mutation counts. While comparing the raw mutation counts to the GFP reporter signal in FACS experiments again yielded remarkable agreement across variants with varying levels of activities, we found that the variance of raw mutation count was not well-calibrated (Figure 3D upper half). We generated a more accurate, calibrated fitness landscape by using a Bayesian denoising model, FLIGHTED, to correct for experimental noise. We trained FLIGHTED for this purpose with raw sequencing reads of each variant, and demonstrated that FLIGHTED can denoise the raw mutation count and generate model-predicted activity of T7 RNA polymerase variants with accurately calibrated errors (Figure 3D lower half). Compared to raw mutation counts, activities generated by FLIGHTED showed better agreement with the GFP reporter signal (Figure 3B) after fitting with a piecewise linear function. We used the denoised data to generate a sequencing logo showing conserved residue amonng the top 50 most active variants (Figure 3C) and three-dimensional fitness landscapes where AA at 756 or 758 are held constant (Figure 3E).

**Figure 3:**
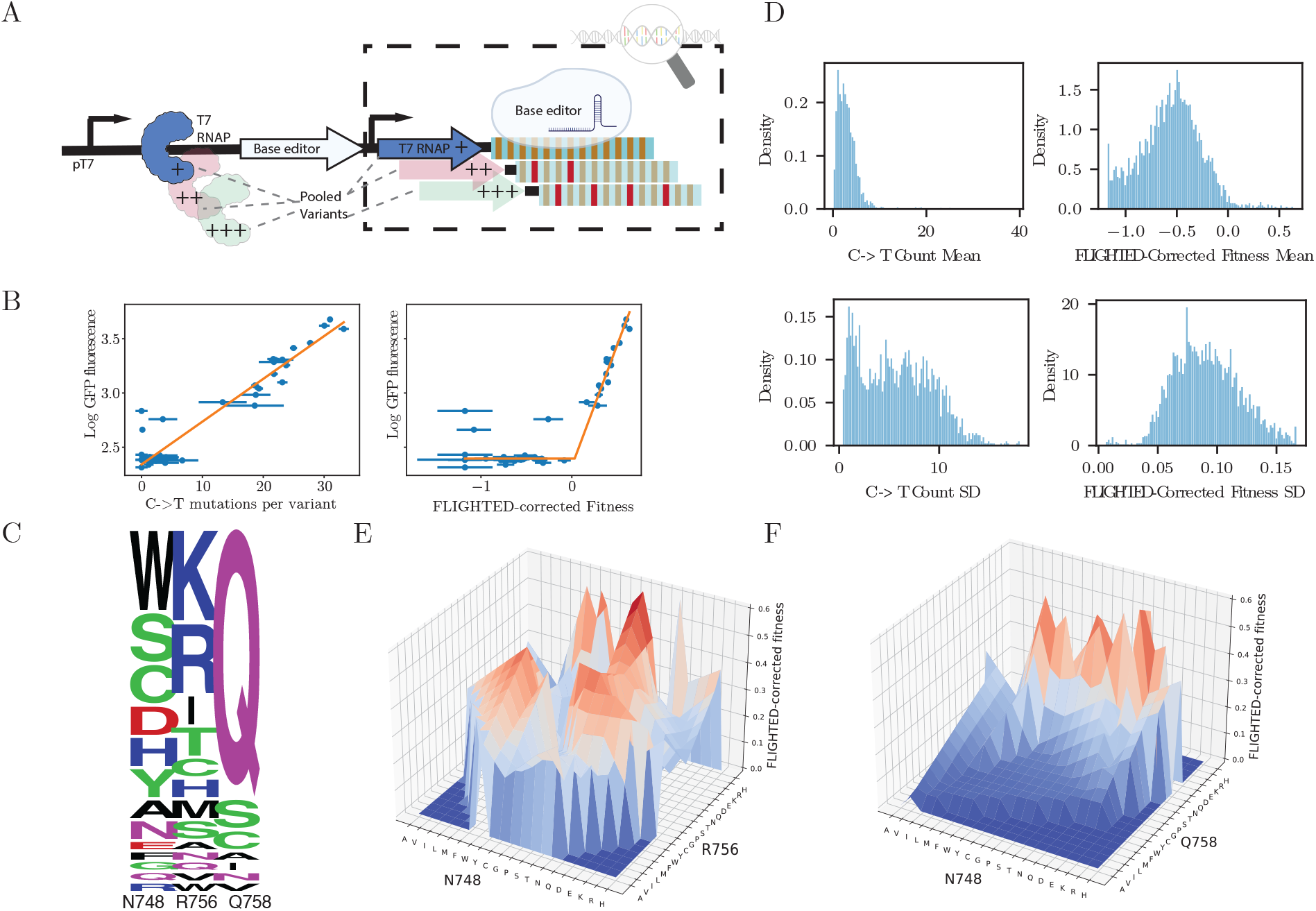
Measuring the activities of roughly 8,000 T7 RNA polymerase variants on T3 promoter with DHARMA. A. Circuit design for T7 RNA polymerase activity measurement. T7 RNAP variants differentially drive the expression of base editor leading to different mutation rate in canvas B. Compared to raw mutation counts, FLIGHTED-corrected fitness show much lower error and better correlation among active variants. C. Sequence logo of T7 RNA polymerase variants. Top 50 most active variants were plotted based on frequency. D. FLIGHTED yields reasonably well-calibrated errors on fitness measurements. E. Fitness landscape of T7 RNAP variants with Q758 held constant. F. Fitness landscape of T7 RNAP variants with R756 held constant.

The three residues we targeted for mutagenesis are part of the specificity loop and known to form hydrogen bonds with the promoter -8 to -11 region [6]. The sequence logo shows that Q758, which forms hydrogen bonds with the adenosine at -8 position, is highly conserved within active variants, while the other two residues, especially N748 which interacts with the promoter at -11 position, are more tolerant to mutations. This is consistent with the sequence identity of the T3 promoter, which differs from the T7 promoter at -11 position but not -8 to -10 positions. While AAs with positively charged side chains such as lysine or arginine are highly preferred at position 756 among active variants, position 748 appears to be tolerant to a wide range of size chain properties, suggesting that promiscuity at position -11 might contribute to increased activity of T7 RNAP on T3 promoter.

### Reconstruct a local fitness landscape of TEV protease with over 100,000 variants

After validating the feasibility of using DHARMA to measure the enzymatic activity of a small library of protein, we shifted focus to quantifying the activity of proteases, a different class of enzyme, by developing a more complex biological circuit. Similar to the T7 RNAP circuit design, the TEV protease circuit involves the transcriptional control of a T7-promoter-driven base editor, the expression level of which is coupled to the activity of TEV protease variant, which cleaves T7 lysozyme, a T7 RNAP inhibitor, both at the N terminus of the T7-RNAP-lysozyme fusion protein, and between the two domains of T7 lysozyme, leading to derepression of base editor transcription. Using the optimized version of this novel biological circuit, we profiled the enzymatic activity of more than 100,000 variants of TEV protease on its wild type substrate and after experimental error calibration using FLIGHTED (Figure 3D), generated the largest TEV protease fitness dataset to date. We performed site-saturation mutagenesis on 4 AA residues (T146, D148, H167, and S170) in the TEV protease S1 pocket, which is known to interact with P1 residue of the substrate and determines substrate specificity. We found that the 99.8% of the library members have fitness values less than that of the wild-type variant (Figure 4C). This is in line with our expectation given that the wild-type sequence has been optimized for activity on the wild-type substrate through natural evolution and any mutation in this key substrate binding pocket is likely to reduce its fitness. Among top 100 most active variants in this dataset, wild-type AA residues (TDHS) or those with similar physical properties are over-represented at each mutable position (Figure 4B). Notably, H167 is the only AA residue that is conserved among all top 100 variants. This is consistent with previous findings that H167 is critical in forming hydrogen bond with the wild-type substrate [7]. While fitness landscape of TEV protease has been previously studied by characterizing a small number of variants generated by directed evolution [8], our large-scale dataset of over 100,000 variants enabled reconstruction of a near-complete 4-site local fitness landscape.

**Figure 4:**
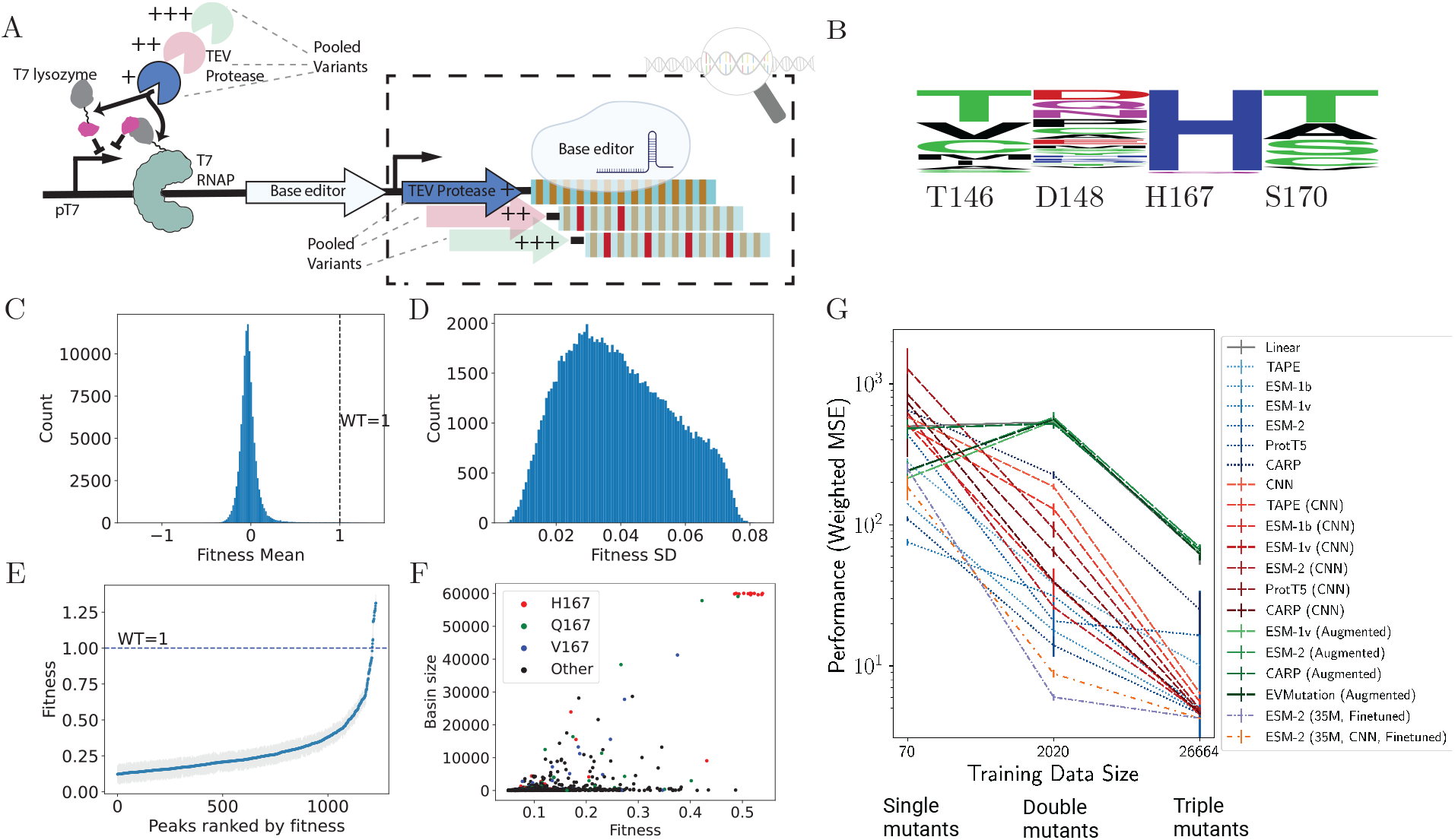
DHARMA enables high-throughput measurement of over 100,000 TEV protease variants in a single experiment. A. Circuit design for TEV protease activity measurement. TEV protease variants differentially derepress the transcription of base editor driven by a T7 RNAP leading to different mutation rates in canvas. B. Sequence logo of top 100 most active TEV protease variants. C. Distribution of fitness values of TEV protease variants. Wild-type variant is scaled to a fitness of 1. Only variants with more than 100 reads are included D. Experimental errors as measured by FLIGHTED. Only variants with more than 100 reads are included. E. Fitness peaks of TEV protease variants. F. Basin sizes of fitness peaks of TEV protease variants. Basin size is defined as the number of variants that are connected to a fitness peak by a directed path. G. Models trained with increasingly larger data are exponentially more accurate.

To identify fitness peaks and assess the ruggedness of the fitness landscape, we first identified all neighboring pairs (one AA mutation away) with at least one functional variant in the pair and generated a directed graph where each node represents a variant and each edge represents a single AA mutation, the direction of which is from the less fit variant to the more fit variant. The largest unidirectionally interconnected part of this directed graph was then extracted for use in subsequent analysis. While over 1,200 fitness peaks (nodes the fitness of which is higher than that of all neighbors) were identified, most of these peaks have fitness well below wild-type level. Nonetheless, the number of fitness peaks suggests that this local TEV protease fitness landscape is highly rugged (Figure 4E). Intriguingly, fitness peaks with higher fitness values tend to have lower variance, which is likely due to the increase in editing event sampling frequency in cells with more baseeditor. This is in contrast with other conventional methods such as phage display [9] where the measurement uncertainty increases with fitness. Therefore, DHARMA potentially allows better characterization of highly active variants, which are often of greater interest in protein engineering.

To characterize the accessibility of fitness peaks, we calculated the number of variants in each basin of attraction, which is defined as a set of variants where a random walk starting from any variant in the set will eventually reach the fitness peak. While this fitness landscape is highly rugged, peaks with high fitness are highly accessible, with basin sizes over 50,000 (Figure 4F). Interestingly, while most high-fitness peaks with H167 are highly accessible, a few highly active peaks with AAs other than histidine are not. This suggests that DHARMA may be used to identify variants with high activity that are not accessible by conventional directed evolution. Taken together, these results demonstrate that DHARMA is a powerful method for highthroughput activity profiling of large libraries and can be used to comprehensively characterize the fitness landscape of proteases.

In the last decade, significant progress has been made in the field of machine learning-based protein engineering [10, 11, 12]. Nonetheless, protein fitness prediction remains a very difficult problem [13, 14]. State-of-the-art protein fitness prediction models such as TranceptionEVE [15], GEMME [16] and MSA Transformer [17], largely rely on unsupervised ML as labeled protein fitness data are scarce – only a total of two million variants have been screened so far across more than 90 experiments in the largest collection of deep mutation scanning data to date [18]. To test whether increasing the amount of training data would improve protein fitness prediction using supervised models, we split the data based on the number of mutations in the coding sequence into single mutants, double mutants and triple mutants and evaluated models trained on these separate datasets which have 70, 2020 and 26664 data points, respectively. As shown in Figure 4G, for the vast majority of models evaluated, the weighted mean square errors (MSE) of model predictions decreases as the amount of training data increases, with a reduction in MSE of more than 2 orders magnitude for the best model. Top-performing models have a weighted MSE of roughly 4 when trained with all triple mutants, i.e. their error on a generic data point is roughly 2x the experimental error. Our findings demonstrate that data size is the single most important factor in determining model performance.

## Discussion

We have developed Direct High-throughput Activity Recording and Measurement Assay (DHARMA), a sequencing-based system for high-throughput characterization of DNA-encoded biomolecular activities with many advantages over existing mainstream methods such as flow cytometry. Based on the principle of molecular recording, DHARMA writes a log of activity history onto a segment of DNA (“canvas”) contiguous to the activity-encoding sequence, and this information is later read via Nanopore sequencing and decoded using a bioinformatics algorithm. The physical linkage between the activity-encoding sequenceand canvas creates a reliable mapping between the identity of the functional sequence and the associated activity in a specific biochemical context, enabling highly multiplexed measurement of DNA-encoded activities in a single Nanopore sequencing run. A large number of diverse DNA sequences can be pooled and directly used input for activity measurement without the need of sequential loading of samples as required for accurate and precise measurement using methods such as FACS. Furthermore, given sufficient read depth, DHARMA provides definitive activity information of every individual variant in the library as opposed to multiple distributions (“bins”) of activities resulting from a group of functionally similar variants, which require subsequent sequencing for identification. Compared to “binning”, measurement on single-variant level is a superior representation of true activity, in particular when the dynamic range of activities is high and the number of “bins” are physically limited.

In future efforts, we aim to enhance our TEV protease circuit to generate datasets with a wider diversity in mutations, extending the DHARMA to a comprehensive library of substrates beyond just the wild-type. Such library-on-library experiments have not been reported in the literature to our best knowledge. We plan to use optimization algorithms to iteratively refine our designs, aiming to improve both the catalytic efficiency and substrate specificity of TEV protease. Engineering TEV protease specificity is challenging using conventional directed evolution methods, which generally struggle to effectively select against undesired activities. Consequently, engineered variants with altered substrate specificity also exhibited pronounced promiscuity [8, 19]. DHARMA enables profiling of an extensive library of variants with ultra-high throughput, systematically identifying sequence features that contribute to both desired and undesired activities. Through optimization and iteration, our goal is to engineer variants that exhibit altered specificity with minimal promiscuity.

Significant advancements have been made in the field of protein machine learning, particularly in structure prediction. However, progress in protein fitness prediction has lagged, with most existing models demonstrating limited effectiveness. These models are predominantly unsupervised due to a scarcity of training data, whereas supervised models typically exhibit superior performance. To address the challenge of data generation for training machine learning models, we introduce a innovative platform, which connects protein fitness or biomolecular activity with the mutation rate of a DNA segment adjacent to the coding sequence of such activities. This sequencing-only quantification approach eliminates the need for costly equipment and has the potential to foster more inclusive, community-driven data collection efforts for protein function prediction and to broadly advance the field of protein engineering.

## Methods

### Plasmid construction

Multilevel Golden Gate cloning and Gibson cloning were used to construct all plasmids used in the experiments. Plasmids were assembled from modules each containing a scarless transcription unit insulated by strong terminators. T4 DNA ligase and Type IIS endonucleases, BsaI, BsmBI, SapI and PaqCI (New England Biolabs, Ipswitch, MA) were used in different levels of modular assembly. To generate a library of plasmids each containing a different promoter, a lacZ*α* cassette containing PaqCI sites was inserted between GFP and base editor RBS sequences via restriction digestion and Gibson cloning. Synthetic dsDNA fragments (Integrated DNA Technologies, Coralville, IA) each containing a different promoter and/or a bidirectional terminator, and a 24-base barcode were inserted scarlessly to replace the lacZ*α* cassette. Blue-white screening was performed as per manufacturer’s instructions. Similarly, to construct a library for circuit optimization, synthetic dsDNA fragments or PCR products containing unique regulatory elements and corresponding barcodes were assembled combinatorially with the coding sequences of interest and the canvas sequence using Golden Gate assembly in a hierarchical manner. To generate library of T7 RNA polymerase variants, site-saturation mutagenesis was performed at AA position 748, 756, and 758 of the T7 RNA polymerase coding sequence using annealed synthetic DNA containing NNK codons and internal barcodes consisting of synonymous codons. These anneal fragments were assembled into a complete gene during library construction at the final level of Golden Gate assembly using a plasmid backbone containing SapI sites and the rest of the components of the circuit. To generate the library of TEV protease variants, site-saturation mutagenesis was performed at AA position 146, 148, 167, and 170 of the TEV protease coding sequence using primers containing NNK codons. The resulting PCR products were assembled into a complete gene during library construction at the final level of Golden Gate assembly using a plasmid backbone containing SapI sites and the rest of the components of the circuit. NEB 10-beta cells (New England Biolabs, Ipswitch, MA) were used for cloning and testing of all constructs. Transformation and selection conditions were based on manufacturer’s recommendations.

### Growth condition and sample collection

After plasmids were successfully cloned, all cells were grown in Davis Rich Media (DRM) [20] to minimize biofilm formation and to lower background noise in fluorescence measurements when applicable. For the promoter library experiment described in Figure 1, commercial 10-beta competent cells (New England Biolabs, Ipswitch, MA) were transformed with plasmids carrying a canvas repeat sequence-targeting sgRNA cassette driven by strong constitutive promoter apFAB36 [21], and rendered electrocompetent using standard procedures. Plasmids containing the base editor were introduced either individually or as a library into the cells above via electroporation at 1700V. Cells were immediately resuspended in SOC medium and allowed to recover at 37°C for 1h. To eliminate plasmids that did not migrate into the cells during electroporation, cells were pelleted, washed with DNaseI reaction buffer (New England Biolabs, Ipswitch, MA), resuspended in DNaseI buffer, and incubated at 37°C for 10min with 2U of DNaseI. Cells were then resuspended in DRM supplemented with appropriate antibiotics at a density of approximately 0.5 OD_600_, and maintained at this density at 37 °C by periodically diluting the cultures with fresh DRM supplemented with appropriate antibiotics. One hundred microliters of cultures were collected hourly for GFP fluorescence measurement and PCR amplification for downstream sequencing. Samples were immediately cooled to 4°C to stop base editing activities. For the T7 RNAP experiment described in Figure 3, electrocompetent cells containing the sgRNA plasmid were prepared and transformed with the T7 RNAP library plasmids as described in the paragraph above. Cells were then resuspended in DRM supplemented with appropriate antibiotics at a density of approximately 0.1 OD_600_, and grown and maintained at a density of 0.5 OD_600_ at 37 °C in a 500mL turbdiostat which monitored the turbidity of the culture and continuously diluted the cultures with fresh DRM supplemented with appropriate antibiotics. Fifty milliliters of samples were taken hourly and immediately cooled to 4°C to stop base editing activities. For the TEV protease experiment described in Figure 4, electrocompetent cells harboring plasmids containing all circuit components except the TEV protease library were prepared and transformed with the TEV protease library plasmids as described in the paragraph above. Cells were then resuspended in DRM, grown and maintained in turbdiostat at 0.5 OD_600_ for 20 hours as described in the paragraph above. Fifty milliliters of samples were taken every two hours and immediately cooled to 4°C to stop base editing activities.

### Reporter assay and Nanopore sequencing

Samples collected as described in the last section were washed with 10mM HEPES buffer and loaded onto a plate reader (BMG Labtech, Ortenberg, Germany) for fluorescence measurement at 470/515 nm (Ex/Em). Absorbance at 600nm was also measured for fluorescence normalization across different cell densities. To amplify the canvas and activity-encoding regions on the recorder plasmid construct,0 5µL of culture sample was directly used in a 10µL PCR reaction using PrimeStar Max master mix (Takara Bio, San Jose, CA) under conditions recommended by the manufacturer. Primers (Azenta Life Sciences, Chelmsford, MA) used in the reactions include 24-base barcodes on the 5′ end to allow highly multiplexed sequencing across different time points on a single flow cell. PCR reactions were pooled and purified with magnetic beads (Aline Biosciences, Woburn, MA) at bead suspension to sample ratio of 0.8-1x. Sequencing library was prepared using the SQK-LSK112 Ligation Sequencing Kit and sequenced on a R10.4 flow cell as per manufacturer’s instructions (Oxford Nanopore Technologies, Oxford, UK).

### Data Analysis

As the activity is recorded in a region contiguous to the functional sequence, it is critical to obtain full-length reads that contain both sequences such that a direct mapping between activity and sequence identity can be established. Nanopore sequencing technology has now been frequently used to generate full-length reads>20kb and the latest chemistry yields a modal raw read accuracy of 99%. Full-length reads of RPs or amplicons of RPs obtained from nanopore sequencing are sufficiently accurate for mutation frequency quantification but present challenge in identifying point mutations or other minor variations within the functional sequence of interest. To this end, we included a 24-base barcode adjacent the sequence of interest during synthesis. These barcodes were used in previous works to demultiplex up to 96 samples with 95% accuracy [22]. For the purpose of system validation, we also performed barcoding PCR directly over the canvas region as an alternative multiplexing method, which was possible in this case because the identity of the functional sequence was known. In the scenario in which the identities of functional sequences cannot be readily associated with an additional identifier, reads can be first grouped by exact matches in the functional sequence region and then further clustered based on error tolerance set for a particular application. These clustered reads are treated as demultiplexed read groups each containing reads with identical functional sequences. Within each read group, the canvas region of each read was aligned to the reference sequence using the Smith–Waterman algorithm [23], and the number of C to T mutations was counted at each position within the canvas region and normalized to the total number of full-length reads in the read group. We generated a mutation profile for each read group and computed the area under the curve as the primary metric of base editing activity. While we observed some degree of noise, potentially caused by sequencing error and non-specific deamination, it was unlikely to materially impact the activity readout due to its low amplitude. These results validated the basic method. For sequencing data generated for the circuit validation experiment using the 24-promoter library, basecalling and demultiplexing were performed using Guppy v5.0.7 (Oxford Nanopore Technologies, Oxford, UK) with the high-accuracy model shipped with the software. Consensus calling was performed within each group of demultiplexed reads using pbdagcon (https://github.com/PacificBiosciences/pbdagcon) to identify or confirm the activity-encoding sequence. Demultiplexed raw reads were then truncated and aligned to the reference sequence of the repetitive canvas region using the Smith-Waterman algorithm implemented in Julia [24]. For each read, the occurrences of mismatch where cytosine was replaced by thymidine and their positions within the canvas region were stored in a binary vector, the index of which corresponds to position relative to the first base of canvas. The sum of these vectors within each demultiplexed read group was normalized with the total number of full-length reads with the group to yield the mutation profile of the canvas sequences associated with a particular library member or variant. The total number of mutations was computed to yield the metric for single-variant level activities. Samples with apparent demultiplexing or amplification problems were excluded. Data smoothing was performed by taking moving average with one neighboring data point. Time series mutation rate data were fitted to a generalized logistic function using the Levenberg–Marquardt algorithm. To validate system performance against independently measured activities, log-transformed total C-to-T mutations for each variant was plotted against corresponding log-transformed fluorescence intensity. Linear regression was performed on the log-transformed data using the least-squares algorithm. For sequencing data generated for the T7 RNAP experiment, basecalling and demultiplexing were performed using Guppy v6.4 (Oxford Nanopore Technologies, Oxford, UK) with the super-high-accuracy model shipped with the software. Individual reads were aligned to reference sequences containing distinct internal barcodes and the mutations in the coding sequence were assigned to the corresponding variant based on the maximum alignment score. The mutations in the canvas sequences were recorded in an array indexed by nucleotide position within the canvas for each variant, with 0 respresenting no mutation, 1 representing a C-to-T mutation and 2 representing mutations other than C-to-T mutations. These arrays together with their corresponding variant AA sequences were used to train FLIGHTED to predict the fitness of each variant. For sequencing data generated for the TEV protease experiments, basecalling and demultiplexing were performed using Dorado v0.5.3 (Oxford Nanopore Technologies, Oxford, UK) with the super-high-accuracy model shipped with the software. Given the high single-molecule accuracy of the latest nanopore chemistry, raw reads were directly used to identify the AA-level mutations in the coding sequence region targeted for site-saturation mutagenesis. Sequences adjacent to the targeted region were located by pairwise alignment against reference sequences and the targeted region was then located and translated to AA sequences. Raw reads with more than 1 mismatch or indel in the targeted region were discarded. The mutations in the canvas sequence were identified and recorded in the same way as described in the T7 RNAP experiment. These arrays together with their corresponding variant AA sequences were used in a pretrained FLIGHTED model to predict the fitness of each variant.

### Software

All sequencing data was processed using custom scripts to be released as part of the DHARMA software package on Github. FLIGHTED software package and the pretrained model used in this study are available at https://github.com/vikram-sundar/FLIGHTED_public.

